# Strategies for effectively modelling promoter-driven gene expression using transfer learning

**DOI:** 10.1101/2023.02.24.529941

**Authors:** Aniketh Janardhan Reddy, Michael H. Herschl, Xinyang Geng, Sathvik Kolli, Amy X. Lu, Aviral Kumar, Patrick D. Hsu, Sergey Levine, Nilah M. Ioannidis

## Abstract

The ability to deliver genetic cargo to human cells is enabling rapid progress in molecular medicine, but designing this cargo for precise expression in specific cell types is a major challenge. Expression is driven by regulatory DNA sequences within short synthetic promoters, but relatively few of these promoters are cell-type-specific. The ability to design cell-type-specific promoters using model-based optimization would be impactful for research and therapeutic applications. However, models of expression from short synthetic promoters (promoter-driven expression) are lacking for most cell types due to insufficient training data in those cell types. Although there are many large datasets of both endogenous expression and promoter-driven expression in other cell types, which provide information that could be used for transfer learning, transfer strategies remain largely unexplored for predicting promoter-driven expression. Here, we propose a variety of pretraining tasks, transfer strategies, and model architectures for modelling promoter-driven expression. To thoroughly evaluate various methods, we propose two benchmarks that reflect data-constrained and large dataset settings. In the data-constrained setting, we find that pretraining followed by transfer learning is highly effective, improving performance by 24 − 27%. In the large dataset setting, transfer learning leads to more modest gains, improving performance by up to 2%. We also propose the best architecture to model promoter-driven expression when training from scratch. The methods we identify are broadly applicable for modelling promoter-driven expression in understudied cell types, and our findings will guide the choice of models that are best suited to designing promoters for gene delivery applications using model-based optimization. Our code and data are available at https://github.com/anikethjr/promoter_models.

## 1. Introduction

Gene therapy aims to deliver therapeutic genes, or transgenes, to disease-associated cells and tissues. The expression of transgenes is controlled by an upstream compact regulatory DNA sequence called a promoter, which consists of transcription factor (TF) binding sites that regulate transcription of the adjacent transgene. To effectively treat disease while mitigating off-target side effects, promoters for gene delivery should be optimized for inducing expression only in particular target cell types (i.e. for inducing differential expression), which requires compact promoter sequences with a high density of regulatory information. Recent advances in single-cell sequencing have illuminated over 400 cell types in the human body (Tabula Sapiens Consortium, 2022), yet only a handful of compact cell-type-specific promoters are known. Traditional methods to engineer promoters with cell-type-specificity rely on manual curation of sequence elements that are known to regulate expression, such as tiling of cis-regulatory elements (CREs) or tandem repeats of TF-binding motifs (Miao et al., 2000; Selvakumaran et al., 2001; Yun et al., 2008; Nissim et al., 2017; Wu et al., 2019). While these approaches have been successful in some cell types, extending them to less-studied cell types is a laborious process.

Data-driven promoter or CRE design methods that harness the power of machine learning (ML) models have been proposed (Linder et al. (2020); Wang et al. (2020); Jores et al. (2021); LaFleur et al. (2022); Gosai et al. (2023) among others). These methods build *sequence-based models* of promoter-driven expression (PE) using experimental measurements and then optimize for the promoter sequence using model predictions as surrogates for experimental measurements. Although these methods have the potential to accelerate promoter discovery by being automated, the models they use are trained on large PE datasets, which are only available for a handful of well-studied cell types, again making it difficult to design promoters that target the vast majority of relatively understudied cell types for which we have relatively small datasets. Additionally, previous promoter design studies have not rigorously explored model architectures and modelling strategies, or explored leveraging existing related datasets for transfer learning to cell types with small datasets, which has the potential to further improve prediction performance. In this work, we aim to address these shortcomings and identify the most effective methods to model PE. We identify and benchmark various architectures and modelling strategies on two PE datasets and show that transfer learning enables us to build accurate sequence-based models of PE in a data-efficient manner, thereby enabling the development and usage of data-driven promoter design methods in data-constrained settings.

Transfer learning using pretrained models has emerged as one of the most effective ways to model small datasets. For example, self-supervised tasks such as masked language modelling (MLM) have recently been used to pretrain genomic sequence embeddings that are then fine-tuned for downstream tasks (e.g. Ji et al. (2021); Mo et al. (2021); Benegas et al. (2022); Zeng et al. (2023)). Pretraining using task-relevant data can improve the performance of fine-tuned models (Gururangan et al., 2020), while pretraining using irrelevant data can hurt performance (Liu et al., 2022). For our application, many datasets are closely related to PE and are potentially useful for transfer learning. In particular, massively parallel reporter assays (MPRAs) typically measure PE from a large set of sequences and this data can be used for model training (e.g. Movva et al. (2019); Agarwal et al. (2023); Gosai et al. (2023)). Data from endogenous sequences have also been used to train large models to predict endogenous gene expression and other molecular phenotypes (Agarwal and Shendure, 2020; Avsec et al., 2021), and these models can be fine-tuned to predict PE (Agarwal et al., 2023). Transcription factor (TF) binding data may also help models learn relevant sequence motifs that regulate expression when present in promoters. We evaluate the utility of pretraining on such datasets for modelling a new small PE dataset that we collected from three immune cell lines, two of which are relatively understudied. We also evaluate the utility of pretraining before fine-tuning on a larger MPRA dataset from five cell lines to understand whether pretraining also helps in this setting.

Our work has three main contributions. Most importantly, we propose and evaluate several transfer learning approaches to improve our ability to model PE and present conclusive evidence that transfer learning significantly improves our ability to predict PE in target cell types, especially in data-constrained settings. As part of this work, we also develop two benchmarks to gauge the performance of PE predictors in both data-constrained and large dataset settings. Finally, we propose a novel model architecture called MTLucifer to effectively model PE datasets when training models from scratch. To the best of our knowledge, prior work has not attempted to use transfer learning to improve PE prediction, apart from Agarwal et al. (2023) who propose to predict PE by performing linear probing on a large model (Enformer) that was previously trained on a variety of endogenous expression and epigenomic data (Avsec et al., 2021). Moreover, unlike prior work that mostly foregoes benchmarking, we systematically benchmark several model architectures and transfer learning methods to identify the best approaches. This benchmarking is performed using two PE datasets: a smaller dataset with ∼ 17*K* PE measurements from three cell lines (**data-constrained setting**), and a larger dataset with ∼ 750*K* PE measurements from five cell lines (**large dataset setting**). In both settings, when models are trained using only the bench-marking datasets (no transfer learning), MTLucifer models have the best performance. When using transfer learning, in the data-constrained setting, we find that Agarwal et al. (2023)’s approach of performing linear probing on Enformer improves prediction performance by **24** − **27**% in all three cell types. We also identify a more inexpensive pretraining approach that pretrains MTLucifer on existing PE data from MPRAs, which improves prediction performance by **10** − **16**%. However, these large performance improvements from pretrained models do not carry over to the large dataset setting, where the best performing method—pretraining MTLucifer on another MPRA dataset before fine-tuning it on the benchmarking dataset—leads to only relatively modest improvements of up to 2% when compared to training models on the target dataset alone. Our findings are broadly applicable to most PE datasets and can be applied to design promoters for gene therapy that are optimized for expression in a therapeutic target cell type, reducing potential off-target side effects in other cell types. Our benchmarks can also be used to thoroughly evaluate the effectiveness of PE predictors.

## 2. Existing gene expression predictors

Endogenous gene expression is a complex process that is regulated by multiple DNA sequence features, including CREs, TF-binding motifs, and epigenetic modifications. Before the advent of DL, most sequence-to-expression models extracted hand-crafted sequence features such as counts of known TF-binding motifs and other short sequence (k-mer) counts within the input sequence (Zrimec et al., 2021). Early applications of DL in genomics used convolutional neural nets (CNNs) with one-hot encoded sequence inputs. For example, Zhou and Troyanskaya (2015) used CNNs to predict various epigenetic modifications and TF-binding sites. More recently, Avsec et al. (2021) showed that using convolutional layers followed by transformer layers (CNN+Transformer model), in a model called Enformer, improves prediction of endogenous gene expression when compared to convolutional layers alone. Although many of these models achieve high accuracy for endogenous expression, they are not suited to directly predicting expression from compact promoters used in gene delivery applications because **(1)** unlike endogenous gene expression, control of promoter-driven expression relies on only a short promoter sequence without additional distal regulatory elements, and **(2)** promoter-driven expression utilizes promoter sequences with a much higher information density (density of regulatory sequence motifs) when compared to endogenous promoters. However, Agarwal et al. (2023) showed that one can build an effective sequence-based PE predictor by training a Lasso regression model (Tibshirani, 1996) that takes Enformer predictions as inputs, using MPRA data. This predictor outperforms two other models that they trained from scratch and shows that models like Enformer encode information that is important for transfer learning since they are typically trained using very large genomic datasets that capture a lot of regulatory grammar.

Models of PE trained exclusively using large MPRA datasets have also been developed, which are more directly relevant to the gene delivery setting. For instance, Movva et al. (2019) train a CNN to predict PE in K-562 and HepG2 cells. Similarly, Gosai et al. (2023) build a CNN to predict PE in K-562, HepG2, and SK-N-SH cells. Recently, the DREAM challenge^1^ aimed to uncover the best architectures to model PE in yeast using a very large MPRA dataset - the best-performing model was a modified CNN called LegNet (Penzar et al., 2022). Although this challenge benchmarked many architectures, transfer learning was not allowed, it did not identify approaches that are ideal for data-constrained settings, and it did not model human cell lines-based MPRA data that is most similar to the gene delivery setting. Therefore, there is a need to identify the best approach to model PE for use in promoter design, including transfer learning approaches, especially in data-constrained settings. To this end, we benchmark many different architectures and see that MTLucifer, a CNN+Transformer model, has the best performance when trained from scratch in both data-constrained and large dataset settings. We then use this architecture, Enformer, and DNABERT (Ji et al., 2021) (a language model trained using the human genome) to explore various transfer learning approaches.

## 3. Transfer learning approaches for leveraging related data

Collecting large datasets that measure PE in multiple cell types is expensive and time-consuming. However, there are several large datasets that provide relevant information for modelling PE, described in Section 6 below. Transfer learning can effectively model small datasets in these settings by leveraging large relevant datasets. In this work, we explore two main types of transfer learning for the PE prediction task: pretraining followed by linear probing or fine-tuning, and joint training. Here, we explain these techniques.

### 3.1. Pretraining followed by linear probing or fine-tuning

When DL models are trained from scratch on small datasets, it is difficult for them to learn all task-relevant features, leading to poor performance. However, if there is a large related dataset, training on that dataset prior to training on the small dataset can help the model learn relevant features that are similar between the two datasets. This procedure is called pretraining. The pretrained model can then be further trained on the small dataset to learn which of the features learned during pretraining are relevant for the task at hand and and to modify their weights as needed. This process is data-efficient, as the model has learned most relevant features during pretraining, and generally leads to better prediction performance on the small dataset (e.g. Devlin et al. (2018); Chen et al. (2020)).

There are two main transfer methods for training on the small dataset after pretraining: linear probing and fine-tuning. Pretrained models generate an embedding of the input before using this embedding to make predictions for the pretraining task. Linear probing freezes all weights of the pretrained model and adds a trainable linear layer that is trained on the small dataset to make predictions for the downstream task of interest using the input embeddings. This training can be regularized using techniques such as Lasso. Fine-tuning not only adds a trainable output linear layer but also allows the weights of the pretrained model to be updated when training on the small dataset. Fine-tuning typically leads to better predictions, but there are some instances where linear probing is better, such as when the small dataset contains inputs that are out-of-distribution for the pretrained model (Kumar et al., 2022).

### 3.2 Joint training

Another effective method to perform transfer learning is to jointly train a model on multiple related datasets, some which are much larger than the target task. Joint training can be accomplished by having a shared backbone network that outputs embeddings of the inputs. These embeddings are then supplied to task-specific layers that output predictions for all tasks. The motivation behind this approach is that the shared backbone network learns a wide variety of features based on the larger datasets, and these features can then be efficiently utilized by the task-specific layers, even for tasks with small training datasets. This method has also been shown to improve prediction performance on the smaller datasets (e.g. Yang et al. (2017)).

#### Performing multi-task learning (MTL)

MTL is required to pretrain or jointly train on multiple tasks. We perform MTL using the torchmtl package (Bock, 2020). A common backbone network is used to embed inputs. The embeddings are then supplied to task-specific linear layers that make task-specific predictions. During training, each batch is composed of samples for one task and we cycle through the tasks while sampling batches in an epoch (batch-level round-robin) which has been shown to be effective (Alayrac et al., 2022). Since the losses for each task can be on different scales, we use Kendall et al. (2018)’s method to learn weightings for each task’s loss. The weighted sum of losses is then minimized using an optimizer.

## 4. Promoter-driven expression datasets used for benchmarking

To evaluate the approaches described in the previous section for training effective PE predictors that leverage large related datasets using transfer learning, we construct two benchmarking datasets. Although we are primarily interested in the more natural data-constrained setting where we have a small target PE dataset, we also wish to evaluate various approaches in the large dataset setting to determine if transfer learning is beneficial in this setting. Thus, we use the two PE datasets described in this section for bench-marking - a small one with ∼ 17*K* measurements that we collected from three cell lines, and a large one with ∼ 750*K* measurements from an existing MPRA performed in five cell lines. Models trained using various strategies are ultimately evaluated in terms of their effectiveness in modelling these two datasets - they simultaneously predict PE (averaged across replicates) in each cell line from the promoter using different output heads.

### 4.1 Fluorescence dataset: small dataset quantifying PE by measuring induced fluorescence levels in a pooled screen (data-constrained setting)

We collect a new relatively small PE dataset from 3 immune cell lines: Jurkat, K-562, and THP-1. These specific cell lines are chosen because of their similarity to primary cells, and because promoters designed for these cell types could be useful for treating blood cancers. Although PE is well-studied in K-562 cells, with multiple MPRAs using K-562s (e.g. Ernst et al. (2016); van Arensbergen et al. (2019)), there are no large scale datasets that measure PE in Jurkats and THP-1s. Thus, we use a pooled screen to measure expression from a set of 20,000 promoters of length 250 base pairs (bp), limited by synthesis constraints similar to a gene therapy setting. We choose our tested promoter sequences using heuristics designed to maximize the number of differentially expressed promoters. Briefly, ∼ 50% of the tested promoters are derived from promoters of differentially expressed endogenous genes (Class I). Another ∼ 40% are designed by tiling known and de-novo motifs that were discovered to be enriched in the promoters of differentially expressed endogenous genes by HOMER (Heinz et al., 2010), a motif detection tool (Class II). The final ∼ 10% of promoters are derived from promoters of highly expressed endogenous genes so that our models can learn features of sequences that lead to high expression across many cell types (Class III).

Each promoter is cloned upstream of a minimal cytomegalovirus (CMV) promoter and the enhanced green fluorescent protein (EGFP) reporter gene into a lentiviral vector. The expression induced in each cell line upon transduction is measured by the induced fluorescence levels, and we collect two replicate measurements of fluorescence. We get adequate data from 17,104 promoters. For model training and evalutation, ∼ 70% of these promoters are included in the training set, ∼ 10% in the validation set, and ∼ 20% in the test set. The promoters in each set are stratified by both promoter class and GC content. More details about the experimental protocol (including how we quantify expression strength) and promoter selection are in Appendix A and B, respectively.

### 4.2. Malinois MPRA: large PE dataset derived from an existing MPRA (large dataset setting)

We use MPRA data from ENCODE (Gosai et al., 2023; ENCODE Project Consortium, 2012) to create a large PE dataset (Appendix C contains ENCODE accession numbers for this data). This data was collected by a single lab using a uniform experimental protocol from five cell lines: GM12878, K562, HepG2, SK-N-SH, and A549. We choose to use this data as it contains a large number of high fidelity measurements from a relatively large number of cell lines. Moreover, a subset of this data has already been used by Gosai et al. (2023) to train a PE predictor called Malinois (hence, we refer to this dataset as Malinois MPRA). The MPRA measures PE from constructs containing 200bp long promoters that are cloned upstream of a reporter gene and delivered to the aforementioned cell lines using transient transfection. PE is roughly computed as the 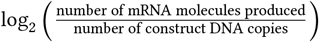. The promoters are mostly human genomic segments containing either the reference or alternate alleles for genomic variants identified by UK Biobank (Sudlow et al., 2015) and GTEx (GTEx Consortium, 2020). We extract 318734, 636185, 750298, 750084, and 318734 PE measurements from GM12878, K562, HepG2, SK-N-SH, and A549 cells respectively. Like Gosai et al. (2023), we use sequences from chromosomes 7, and 13 for testing (∼ 13% of all sequences), those from chromosomes 19, 21, and X for validation (∼ 7% of all sequences), and all other sequences for training.

## 5. Model architectures for benchmarking

We need effective model architectures to make the best use of the available data. Here, we briefly describe the various model architectures that we benchmark and our rationale for choosing to benchmark them. All models, except the motif occurrences-based ones, are sequence-based, and take one-hot encoded sequences as inputs. Models are described in more detail in Appendix E.

### MTLucifer

We propose a smaller CNN+Transformer architecture inspired by Enformer (Avsec et al., 2021) called MTLucifer. Since promoters are relatively short sequences, models such as Enformer that use pooling layers in their CNNs could lose granular information that might be important for modelling promoters. Therefore, we choose to use 3 length-preserving convolutional layers followed by 5 transformer layers in MTLucifer. A [CLS] token embedding is appended before the transformer layers and its final embedding is used by the output layers to make predictions.

### Motif occurrences-based fully connected networks (FCN)

To compare sequence-based modelling strategies with more conventional strategies that represent a sequence using handcrafted features, we evaluate two FCNs (one with 4 layers and a larger one with 6) that take vectors of known TF-binding motif occurrences in the promoters as inputs.

### CNNs

To evaluate if transformer layers are beneficial for modelling PE, we benchmark 3 CNNs. The first CNN uses 4 convolutional layers while the second larger CNN uses 6 such layers. The last CNN is a ResNet with 8 residual blocks. In all 3 CNNs, the outputs of the convolutional layers are supplied to fully connected layers that make predictions.

The next three sets of models are derived from recent work that also aimed to predict PE. We include these models to evaluate their performance on the benchmark datasets.

### LegNets (Penzar et al., 2022)

As mentioned in Section 2, LegNets were the best predictors of PE in yeast in the DREAM challenge. We benchmark two LegNets - one with the same structure as the model that won the challenge, and a larger one with more filters in every convolutional layer.

### MPRAnn (Agarwal et al., 2023)

MPRAnn is a recently proposed 4-layer CNN for modelling MPRA data.

### Malinois (Gosai et al., 2023)

Malinois is a 3-layer CNN that was used to model a portion of the Malinois MPRA dataset that we use for benchmarking.

### DNABERT (Ji et al., 2021)

DNABERT is a BERT model (Devlin et al., 2018) trained using the human genome. To evaluate whether MLM-based pretraining helps in modelling PE, we finetune DNABERT on our benchmark datasets and evaluate its performance.

### Enformer (Avsec et al., 2021)

Enformer is a powerful CNN+Transformer-based gene expression predictor trained using a large set of genomic and epigenomic data, including endogenous gene expression data. Agarwal et al. (2023) showed that Enformer can be used to accurately model PE. We first benchmark a randomly initialized Enformer model. Then, we perform finetuning and linear probing on a pretrained model. This allows us to simultaneously study the merits of the architecture, and the effect of pretraining.

## 6. Pretraining or joint training tasks

In the previous sections, we described the transfer learning methods we adopt, benchmark datasets, and model architectures. DNABERT and Enformer are trained using MLM and a large genomic dataset respectively. Here, we identify four additional large relevant genomic datasets that can be used for pretraining or joint training. Crucially, these datasets are much smaller than the datasets used to train DNABERT and Enformer, making it feasible to perform pretraining on a limited compute budget. This flexibility could allow us to easily explore alternate architectures, hyperparameters, and modelling frameworks. In our experiments, we train MTLucifer models on these datasets to determine their usefulness for transfer learning. More details about some tasks are in Appendix D and Supplementary Table S.1 summarizes them.

### RNA-sequencing (RNA-Seq) data

As endogenous promoters play a crucial role in gene expression, it might be useful to pretrain models on endogenous expression data measured using RNA-seq in various cell types. This should enable the model to learn TF-binding motifs and their relative importances in various cell types. Thus, we pretrain on three large RNA-Seq datasets: LL-100 (Quentmeier et al., 2019), CCLE (Barretina et al., 2012), and Roadmap (Kundaje et al., 2015). LL-100, CCLE, and Roadmap contain expression values from 100, 1408, and 56 cell lines, respectively. From each dataset, we get expression values in every cell line, as measured by TPM or RPKM values. Then, we extract 250bp promoter regions for every gene to input them to our models and predict expression. Genes from distinct chromosomes are used in the train, test, and validation sets: ∼ 70%, ∼ 20% and ∼ 10% of the overall genes are assigned to the train, test, and validation sets, respectively.

### ENCODE TF-binding ChIP-seq data

ChIP-seq assays are used to discover genomic regions that are bound by TFs, and pretraining on such data can help models learn TF-binding sequence motifs. We obtain ChIP-seq peaks and their corresponding sequences for 1363 cell types from ENCODE. Then, we pretrain our models to predict whether a given sequence contains a peak in each of the 1363 cell types. The positive set for this classification task consists of 600bp sequences centered at every peak. In total, there are ∼ 3M peaks. The negative set is built by sampling a dinucleotide shuffled sequence for every positive sequence, similar to the approach followed by Alipanahi et al. (2015) and Zeng et al. (2016). Peaks (and their corresponding negative sequences) from distinct chromosomes are used in the train, test, and validation sets with ∼ 66.8%, ∼ 23.6%, and ∼ 9.6% of the peaks assigned to the train, test, and validation sets, respectively.

### Sharpr-MPRA data

MPRAs measure promoter-driven expression induced by multiple promoters in parallel and thus have high throughput. We hypothesize that pretraining on MPRA data might be very beneficial for our task because of the similarity in experimental protocols - the main difference being that our data measures expression induced by stable transduction while MPRAs measure expression induced by transient transfection. The Sharpr-MPRA dataset (Ernst et al., 2016) measures expression induced by ∼ 487K 145bp promoters in K-562 and HepG2 cells. These promoters are derived from DNase I peaks in K-562, HepG2, HUVEC, and H1-hESC cells. Each promoter is cloned upstream of a minimal TATA or strong SV40 promoter and promoter-driven expression is measured for both conditions. Two replicates of these measurements are collected. Thus, there are 8 measurements per promoter (2 cell lines, 2 downstream promoters, 2 replicates). This dataset was also modelled by Movva et al. (2019), who include each promoter’s reverse complement as an additional training example with the same associated expression value. They also predict the average of the values from the two replicates, leading to 12 outputs per input sequence. The ∼ 20K sequences from chromosome 18 and the ∼ 30K sequences from chromosome 8 are used for testing and validation, respectively. All other sequences are used for training. We use their processed data and modelling setup for pretraining.

### SuRE MPRA data

SuRE (van Arensbergen et al., 2017) is another MPRA that was scaled up by van Arensbergen et al. (2019) to survey the genomes of 4 individuals from 4 different populations. The genomes of these individuals are broken into 150-500bp fragments and each fragment is cloned into a reporter plasmid. These sequence fragments can drive expression and function as promoters in transfected cells if the fragment contains a valid TSS. ∼ 2.4B and ∼ 1.2B fragments were found to be expressed in K-562 and HepG2 cells, respectively. Pretraining on this large dataset allows our models to learn about the structure of promoters and the effects of single nucleotide polymorphisms (SNPs) on expression.

To the best of our knowledge, no other study has used this data for pretraining. Since pretraining on the full dataset is time-consuming due to its size, we subsample it and create a classification task. Our sub-sampling accounts for GC content to reduce any associated confounding. First, each tested sequence is binned into 2 expression bins, one for K-562 and one for HepG2. We define 5 bins for each cell based on the number of reads associated with each sequence: 0, (0, 10], (10, 20], (20, 30] and 30+. Most sequences have 0 reads and the number of sequences assigned to each bin decreases with higher read counts. We remove any sequences with ambiguous SNPs and compute the GC content of each sequence. For each individual, we compute a histogram of GC content over all sequences from their genome, with a bin width of 0.05. Then, for each individual and for each combination of K-562 and HepG2 expression bins (25 combinations), we subsample the individual’s sequences in that bin combination while keeping the GC content distribution as close as possible to the overall GC content distribution. We aim to get 30K training sequences and 3K testing and validation sequences from each bin combination, reflecting different levels of differential expression; however, some bin combinations have fewer sequences. Ultimately, we obtain ∼ 400 − 600K training sequences per individual and ∼ 50 − 70K testing and validation sequences. We create datasets for each individual separately. Our models are pretrained to predict a sequence’s K-562 and HepG2 expression bin in every individual.

## 7. Results

Here we evaluate the model architectures, transfer learning methods, and pretraining tasks described above using our benchmarking datasets in both the data constrained and large data settings. First, we test the various model architectures by training randomly initialized models from scratch using the benchmarking datasets. Then, we evaluate the efficacy of various transfer learning methods. Finally, we demonstrate the usefulness of our trained models in filtering out promoters with low expression, or low PE, a task that is crucial for efficient promoter design. In all our tables and figures, *r* denotes the Pearson correlation coefficient and *ρ* denotes the Spearman’s rank correlation coefficient between the predictions and targets. Experimental hyperparameters are detailed in Appendix E.

### 7.1 Evaluating model architectures

Before evaluating the benefits of transfer learning, we first evaluate the effectiveness of modelling our two PE datasets using each of the architectures mentioned in Section 5, without any pretraining. Tables 1 and 2 show our results on the fluorescence (data-constrained setting) and Malinois MPRA (large dataset setting) datasets respectively. From the tables, we see that MTLucifer is generally the best performing architecture, producing the most accurate predictions for most cells in both the data-constrained and large dataset settings. The large LegNet is also competitive, especially in the data-constrained setting, and the Malinois model that was proposed to model a subset of the Malinois MPRA dataset produces good predictions for that dataset. We can also conclude that sequence-based models are superior to models that use handcrafted features such as motif occurrence counts. Moreover, using a CNN+Transformer instead of a CNN boosts performance. Finally, the relatively poor performance of a randomly initialized Enformer suggests that its architecture is not naturally suited to model PE.

**Table 1:** Average prediction performance obtained using various model architectures when trained from scratch on the fluorescence dataset. The mean and standard deviation are obtained by fitting 5 different models using 5 different train, test and validation splits of the data.

| Model Class | Number of parameters | Jurkat |  | K-562 |  | THP-1 |  |
| --- | --- | --- | --- | --- | --- | --- | --- |
| | | $r$ | $\rho$ | $r$ | $\rho$ | $r$ | $\rho$ |
| MTLucifer | 66.3M | 0.6129 ± 0.0151 | 0.5713 ± 0.0171 | **0.5991 ± 0.0134** | **0.5844 ± 0.0117** | **0.5556 ± 0.0200** | **0.4745 ± 0.0122** |
| Motif-based FCN | 5.1M | 0.4663 ± 0.0153 | 0.4536 ± 0.0136 | 0.4662 ± 0.0058 | 0.4757 ± 0.0129 | 0.4055 ± 0.0158 | 0.3785 ± 0.0142 |
| Large motif-based FCN | 38.1M | 0.4583 ± 0.0177 | 0.4408 ± 0.0089 | 0.4576 ± 0.0115 | 0.4643 ± 0.0141 | 0.4011 ± 0.0146 | 0.3701 ± 0.0171 |
| CNN | 11.0M | 0.5312 ± 0.0168 | 0.4608 ± 0.0126 | 0.5068 ± 0.0094 | 0.4765 ± 0.0065 | 0.4804 ± 0.0130 | 0.3802 ± 0.0096 |
| Large CNN | 21.5M | 0.5485 ± 0.0106 | 0.4629 ± 0.0098 | 0.5212 ± 0.0123 | 0.4771 ± 0.0102 | 0.5064 ± 0.0170 | 0.3797 ± 0.0107 |
| ResNet | 114M | 0.5243 ± 0.0176 | 0.4672 ± 0.0132 | 0.5097 ± 0.0185 | 0.4842 ± 0.0112 | 0.4716 ± 0.0237 | 0.3955 ± 0.0116 |
| LegNet | 1.8M | 0.5729 ± 0.0164 | 0.5234 ± 0.0128 | 0.5551 ± 0.0158 | 0.5354 ± 0.0127 | 0.5035 ± 0.0215 | 0.4270 ± 0.0164 |
| Large LegNet | 33.1M | **0.6156 ± 0.0071** | **0.5747 ± 0.0124** | 0.5876 ± 0.0176 | 0.5785 ± 0.0094 | 0.5490 ± 0.0174 | 0.4639 ± 0.0086 |
| MPRAnn | 807K | 0.5578 ± 0.0131 | 0.5088 ± 0.0233 | 0.5366 ± 0.0129 | 0.5198 ± 0.0229 | 0.4860 ± 0.0152 | 0.4170 ± 0.0238 |
| Malinois | 4.1M | 0.5025 ± 0.0196 | 0.4798 ± 0.0187 | 0.4900 ± 0.0191 | 0.4918 ± 0.0145 | 0.4338 ± 0.0292 | 0.3844 ± 0.0188 |
| DNABERT (random initialization) | 89.2M | 0.5886 ± 0.0125 | 0.5492 ± 0.0113 | 0.5695 ± 0.0117 | 0.5480 ± 0.0117 | 0.5202 ± 0.0197 | 0.4538 ± 0.0130 |
| Enformer (random initialization) | 229M | 0.5401 ± 0.0157 | 0.4959 ± 0.0339 | 0.5234 ± 0.0236 | 0.5039 ± 0.0271 | 0.5061 ± 0.0270 | 0.4118 ± 0.0355 |
| Test Set Replicate Concordance | | 0.7900 ± 0.0271 | 0.7348 ± 0.0116 | 0.7267 ± 0.0247 | 0.6875 ± 0.0093 | 0.6561 ± 0.0423 | 0.4987 ± 0.0133 |

**Table 2:** Prediction performance obtained using various model architectures when trained from scratch on the Malinois MPRA dataset.

| Model Class | Number of parameters | HepG2 |  | K-562 |  | SK-N-SH |  | A549 |  | GM12878 |  |
| --- | --- | --- | --- | --- | --- | --- | --- | --- | --- | --- | --- |
| | | $r$ | $\rho$ | $r$ | $\rho$ | $r$ | $\rho$ | $r$ | $\rho$ | $r$ | $\rho$ |
| MTLucifer | 66.3M | **0.8102** | **0.7775** | **0.8068** | **0.7513** | **0.8055** | **0.7802** | 0.6919 | 0.6679 | 0.4429 | 0.4618 |
| Motif-based FCN | 5.1M | 0.4039 | 0.2925 | 0.3562 | 0.2244 | 0.3944 | 0.3092 | 0.3409 | 0.2671 | 0.2181 | 0.1725 |
| Large motif-based FCN | 38.1M | 0.3930 | 0.2941 | 0.3462 | 0.2246 | 0.3827 | 0.3094 | 0.3316 | 0.2637 | 0.2036 | 0.1811 |
| CNN | 11.0M | 0.7474 | 0.6959 | 0.7294 | 0.6489 | 0.7543 | 0.7180 | 0.6324 | 0.5826 | 0.3949 | 0.3903 |
| Large CNN | 21.5M | 0.7411 | 0.6895 | 0.7151 | 0.6318 | 0.7492 | 0.7139 | 0.6314 | 0.5926 | 0.3987 | 0.3953 |
| ResNet | 114M | 0.7628 | 0.7114 | 0.7399 | 0.6635 | 0.7672 | 0.7358 | 0.6558 | 0.6140 | 0.3969 | 0.3920 |
| LegNet | 1.8M | 0.7955 | 0.7500 | 0.7772 | 0.7156 | 0.7894 | 0.7561 | 0.6898 | 0.6493 | 0.4402 | 0.4404 |
| Large LegNet | 33.1M | 0.8051 | 0.7595 | 0.7871 | 0.7278 | 0.7972 | 0.7592 | 0.6959 | 0.6571 | 0.4450 | 0.4464 |
| MPRAnn | 808K | 0.5295 | 0.4556 | 0.4436 | 0.3594 | 0.5477 | 0.4870 | 0.3733 | 0.3608 | 0.1873 | 0.2011 |
| Malinois | 4.5M | 0.7935 | 0.7633 | 0.7904 | 0.7401 | 0.7924 | 0.7687 | **0.6973** | **0.6720** | **0.4595** | **0.4882** |
| DNABERT (random initialization) | 89.2M | 0.6516 | 0.6256 | 0.6046 | 0.5710 | 0.6638 | 0.6524 | 0.5327 | 0.5173 | 0.3469 | 0.3432 |
| Enformer (random initialization) | 229M | 0.7152 | 0.6409 | 0.6700 | 0.5765 | 0.7158 | 0.6529 | 0.5779 | 0.5169 | 0.3644 | 0.3646 |

### 7.2. Evaluating transfer learning methods

Next, we systematically evaluate various transfer learning-based training strategies. For evaluating the usefulness of the datasets described in Section 6, we use the MTLucifer architecture since it had the highest overall performance when trained from scratch. We pretrain it using each of the tasks in Section 6, and also using some combinations. Then, we either perform linear probing or fine-tuning to model the benchmark PE datasets. Similarly, we perform joint training by training on the various tasks in addition to the benchmark tasks. To evaluate the usefulness of existing pretrained models, we also perform fine-tuning on Enformer and DNABERT. Specifically for Enformer, to replicate the method proposed by Agarwal et al. (2023), we try linear probing on its outputs using Lasso (Tibshirani, 1996).

Tables 3 and 4 present our results. When modelling the fluorescence data, linear probing of Enformer using Lasso is the best performing method, improving the *ρ* by 24 − 27% when compared to the best model that was trained from scratch. This results highlights the importance of pretraining pretraining Enformer using a large genomic dataset makes it the best PE predictor even though its architecture is not naturally suited for predicting PE, as mentioned previously. We also see that fine-tuning an MTLucifer model pretrained using other MPRA datasets improves *ρ* by 10 − 16%, demonstrating that pretraining on these datasets is also useful for modelling the fluorescence data. Moreover, pretraining on this data takes 33 hours on a single Nvidia A40 GPU and training Enformer takes 3 days on 64 TPU v3 cores (Avsec et al., 2021). Therefore, pretraining on the existing MPRA data is much more compute-efficient and enables us to try different architectures, hyperparameters, and modelling frameworks if necessary, while still being assured of good downstream performance.

**Table 3:**
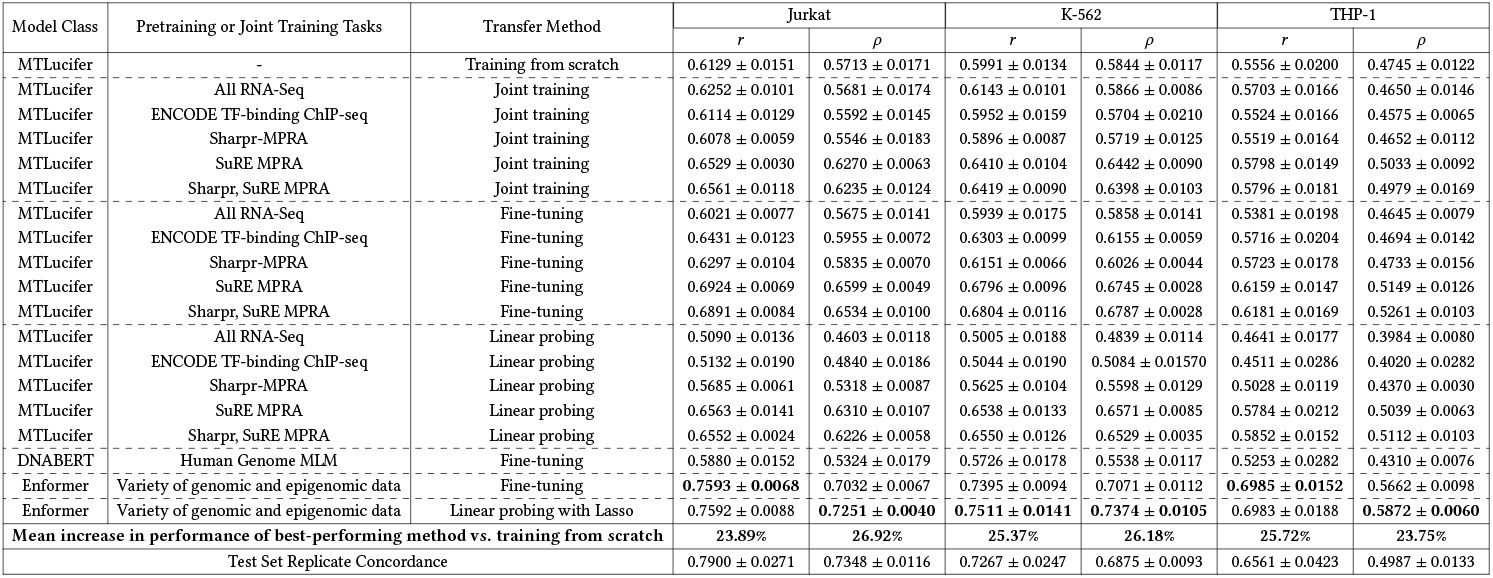
Average prediction performances obtained using various training strategies when used to model the fluorescence dataset. The mean and standard deviation is computed by fitting 5 different models using 5 different train, test and validation splits of the data.

**Table 4:**
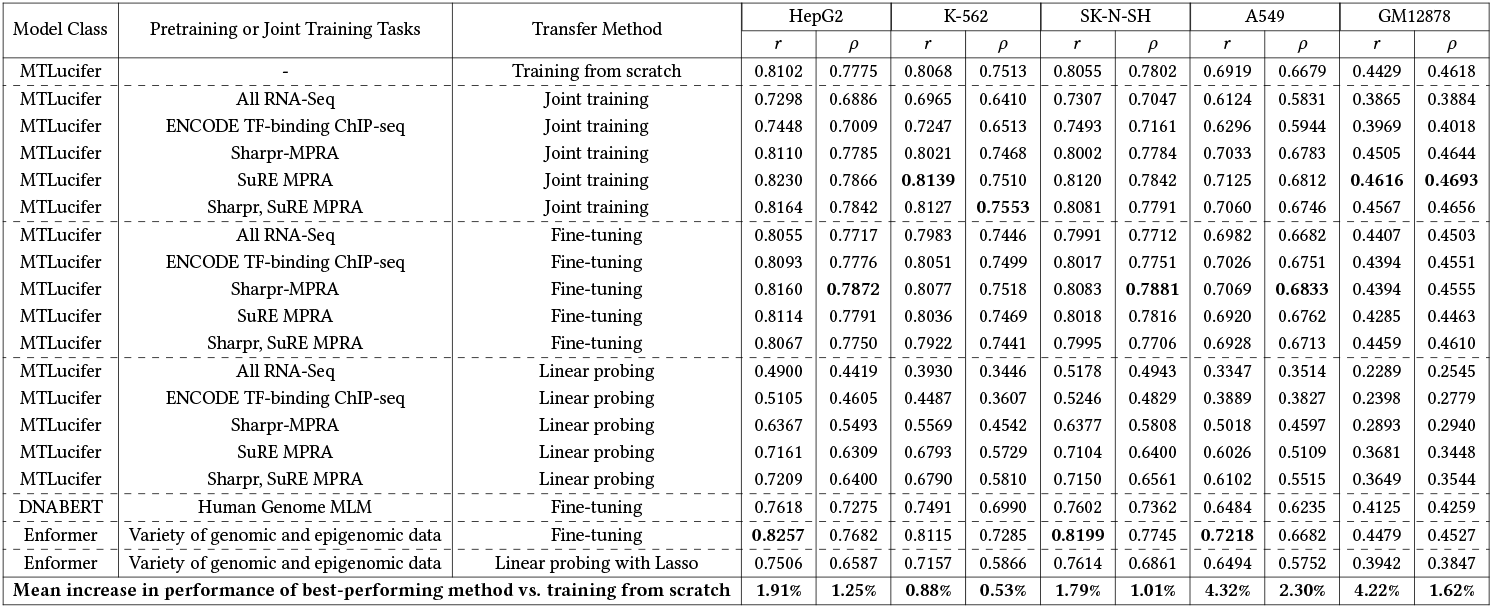
Prediction performances obtained using various training strategies when used to model the Malinois MPRA dataset.

When modelling the Malinois MPRA data and operating in the large dataset setting, we notice more modest performance gains from transfer learning. Here, the best performing method pretrains an MTLucifer model on the Sharpr-MPRA dataset before fine-tuning it on the Malinois MPRA data, and improves the *ρ* by up to 2% compared to training from scratch.

We also notice some instances of negative transfer (performance drops when using a pretrained model vs. a randomly initialized one) such as when we pretrain on RNA-Seq data, highlighting the importance of carefully validating the usefulness of pretraining on a certain dataset.

The smaller fluorescence dataset used to evaluate approaches in the data-constrained setting and the larger Malinois MPRA dataset used to evaluate approaches in the large dataset setting differ in the experimental assay used to collect the data and also in the composition of the tested sequences. To show that our results in the data-constrained setting hold when using either type of data, in Appendix F, we benchmark the various architectures and transfer learning approaches on a subsampled version of the Malinois MPRA dataset whose training set is similar in size to the training set of the fluorescence dataset. We obtain results that are similar to those obtained using the fluorescence dataset, confirming that the size of the training dataset is the main determinant of relative performances.

In conclusion, transfer learning largely improves PE prediction performance but the improvement is more pronounced in the data-constrained setting.

### 7.3. Detecting low PE promoters

Detecting low PE sequences is crucial for efficient promoter design - we want to avoid testing sequences that have low expression in the target cells since they are unsuitable for gene therapies. To determine whether transfer learning helps in detecting such sequences, we create a binary classification task using the fluorescence data - each promoter is assigned three binary labels indicating whether its PE was above the median PE in each of the three cell types. We then build three models to perform this task - an MTLucifer model trained from scratch, a fine-tuned Enformer model, and a fine-tuned MTLucifer model that was pretrained on the SuRE and Sharpr-MPRA data.

Our results are presented in Table 5. The benefits of transfer learning are clear - fine-tuning Enformer improves overall prediction accuracy by 7 −8% when compared to training from scratch. More interestingly, when we analyze highly and lowly expressed promoters for each cell type (defined as the top and bottom 10%iles of promoters, respectively), we find that fine-tuned Enformer significantly increases prediction accuracy on lowly expressed promoters (by 15−21%), while maintaining high accuracy on highly expressed promoters. This indicates that the performance gains obtained by pretraining can greatly improve our ability to filter out lowly expressed promoters.

**Table 5:** Performance of models on the binary classification task constructed using the fluorescence datasets. The mean and standard deviation is computed by fitting 5 different models using 5 different train, test and validation splits of the fluorescence dataset.

| Metric | Performance of MTLucifer model trained from scratch | Performance of fine-tuned MTLucifer model pretrained on SuRE and Sharpr-MPRA data | Performance of fine-tuned Enformer model | Increase in performance of best-performing method vs. training from scratch |
| --- | --- | --- | --- | --- |
| Overall Accuracy Jurkat | 0.7204 ± 0.0147 | 0.7513 ± 0.0117 | **0.7757 ± 0.0095** | 7.68% |
| Overall Accuracy K-562 | 0.7288 ± 0.0085 | 0.7732 ± 0.0070 | **0.7788 ± 0.0039** | 6.86% |
| Overall Accuracy THP-1 | 0.6714 ± 0.0104 | 0.7033 ± 0.0078 | **0.7166 ± 0.0062** | 6.73% |
| Top 10%ile Accuracy Jurkat | 0.9083 ± 0.0319 | 0.9512 ± 0.0114 | **0.9530 ± 0.0109** | 4.92% |
| Top 10%ile Accuracy K-562 | 0.9137 ± 0.0136 | **0.9649 ± 0.0226** | 0.9524 ± 0.0038 | 5.60% |
| Top 10%ile Accuracy THP-1 | 0.8988 ± 0.0437 | **0.9351 ± 0.0099** | 0.9274 ± 0.0144 | 4.04% |
| Bottom 10%ile Accuracy Jurkat | 0.7577 ± 0.0276 | 0.7988 ± 0.0532 | **0.8851 ± 0.0246** | 16.81% |
| Bottom 10%ile Accuracy K-562 | 0.7631 ± 0.0364 | 0.8214 ± 0.0409 | **0.8804 ± 0.0298** | 15.37% |
| Bottom 10%ile Accuracy THP-1 | 0.6530 ± 0.0879 | 0.7065 ± 0.0419 | **0.7964 ± 0.0293** | 21.96% |

## 8. Conclusion

We identify several transfer learning approaches to effectively model PE. We propose two benchmark datasets to analyze the effectiveness of various approaches in modelling PE in data-constrained and large dataset settings. When we evaluate models trained from scratch, MTLucifer, a CNN+Transformer model we propose, generally has the best performance in both settings. Moreover, when we employ transfer learning, we notice significant increases in prediction performance compared to training from scratch. In the data-constrained setting, we see an improvement of 24 − 27%, obtained by performing linear probing on Enformer (Avsec et al., 2021) outputs using Lasso. We also identify a more compute-efficient pretraining approach that improves performance by 10 − 16% - it pretrains an MTLucifer model on SuRE and Sharpr-MPRA data (existing PE data) before fine-tuning it on the benchmark dataset. In the large dataset setting, we see modest gains of up to 2% using the best approach that pretrains an MTLucifer model on the Sharpr-MPRA dataset before fine-tuning it on the benchmarking dataset. Finally, we show the utility of our accurate PE predictors in identifying undesirable low expression promoters - a fine-tuned Enformer model can filter out low expression promoters with 15 − 21% higher accuracy that the best model that was trained from scratch, further highlighting the utility of transfer learning. Our methods and results are useful for modelling any PE dataset, and future work can use our benchmarks to evaluate novel approaches against existing ones.

## Acknowledgments

This work was partially supported by an H2H8 Graduate Research Grant to AJR, a Schmidt Futures grant to SL, and the U.S. National Institutes of Health grant R00HG009677 to NMI.

This research used the Savio computational cluster resource provided by the Berkeley Research Computing program at the University of California, Berkeley (supported by the UC Berkeley Chancellor, Vice Chancellor for Research, and Chief Information Officer).

## A. Experimental methods

### A.1. Library cloning

The promoter library was synthesized by Twist Biosciences in a pooled fashion using microarray sythethesis. 25bp overhangs were added to each 250bp promoter sequence to allow for PCR amplification and Golden Gate assembly (5’-TAGTCGGCTAGATGCGTCTCCTACG(Nx250)GGTACGAGACGACTGTCTTTCCCCT-3’). 20ng of the oligopool was PCR amplified in a 50µL reaction using 1.5µL of of each 10µM primer (TAGTCGGCTAGATGCGTCTCC and AGGGGAAAGACAGTCGTCTCG), and 25µL KAPA HiFi HotStart ReadyMix (Roche KK2602). The thermocycling protocol was 98°C for 3 minutes followed by 12 cycles of 98°C for 20s, 69°C for 15s, 72°C for 15s with a final extension at 72°C for 1 minute. 1µL of the reaction was analyzed by gel electrophoresis, and a single band was visualized at 300bp. The remainder of the reaction was purified using DNA Clean & Concentrator-5 (Zymo D4004) and eluted in 12µL of nuclease-free H_2_O. The amplified oligopool was then cloned into a 3rd generation lentiviral vector immediately upstream of a minimal CMV promoter driving the expression of enhanced green fluorescent protein (EGFP) using a 25µL Golden Gate reaction containing 250ng backbone plasmid, 2X molar of the purified oligopool, 1µL Esp3I (Thermo Fisher FD0454), 1µL T4 DNA ligase (NEB M0202L, 400U/µL) and 2.5µL T4 ligase buffer (NEB B0202S). After an initial 5 minute digestion at 37°C, 30 cycles of 37°C digestion and 16°C ligation were followed by 20 minutes of ligation at 16°C, 30 minutes of digestion at 37°C and 20 minutes of heat-inactivation at 80°C. The reaction was purified using DNA Clean & Concentrator-5 (Zymo D4004) and eluted in 6µL of nuclease-free H_2_O. 2µL were transformed into Endura electrocompetent cells (Biosearch Technologies 60242-2) following the manufacturer’s protocol. After recovery, the cells were plated on a single large 245mm x 245mm LB plate with carbenicillin, and serial dilutions were plated on standard sized plates up to 1:1×10^6^ to assess library coverage. After overnight incubation at 30°C, colonies were counted on the dilution plates to assure a library coverage of at least 30X. Colonies from the large plate were scraped into liquid suspension and collected into a 50mL conical tube before the plasmid pool was prepared using NucleoBond Xtra Midi EF (Macherey-Nagel 740420). Subsequent analysis of the plasmid pool using gel electrophoresis confirmed a homogenously sized plasmid species that was not digestible with Esp3I (Thermo Fisher FD0454).

### A.2. Cell lines and culture conditions

Jurkat, K-562, and THP-1 cells were obtained from American Type Culture Collection (TIB-152, CCL-243, and TIB-202) and grown in RPMI + GlutaMAX (Gibco 61870036) supplemented with 10% FBS (Gibco 26140079), 1x penicillin/streptomycin (Gibco 15140122), 1mM sodium pyruvate (Gibco 11360070) and 10mM HEPES (Gibco 15630080). Jurkat and K-562 cells were generally maintained between 1×10^5^-1×10^6^ cells/mL, and THP-1 wells were maintained between 2×10^5^-1×10^6^ cells/mL. All suspension cell lines were split every 2-4 days by counting cell density and diluting cells into a new flask with fresh medium warmed to 37°C. Lenti-X 293T cells were attained from Takara Bio (632180) and grown in DMEM, high glucose, pyruvate (Gibco 11995065) supplemented with 10% FBS (Gibco 26140079) and 1x penicillin/streptomycin (Gibco 15140122). Lenti-X cells were split every 2-4 days by aspirating medium, treating with TrypLE Express (Gibco 12604021), and reseeding cells into a new flask with fresh medium warmed to 37°C. Incubator conditions were kept at 37°C, 5% CO_2_ and >90% RH. All cell lines were routinely tested for mycoplasma contamination every 2-4 months with MycoStrip mycoplasma detection kit (InvivoGen rep-mysnc-100).

### A.3. Lentiviral production and titration

Large scale lentiviral production was performed in Lenti-X cells by polyethylenimine (PEI, Polysciences 23966) transfection into confluent T225 flasks containing DMEM, high glucose, pyruvate (Gibco 11995065) supplemented with 10% FBS (Gibco 26140079) and 10mM HEPES (Gibco 15630080). 40µg of DNA were transfected into each flask using 2nd generation packaging plasmids pMD2.G (Addgene #12259) and psPAX2 (Addgene #12260) along with the lentiviral plasmid pool at a mass ratio of 1:2:4. After 72 hours of incubation, lentiviral particles were concentrated 10X using Lenti-X Concentrator (Takara Bio 631232) per the manufacturer’s instructions, and single use aliquots were frozen at -80°C. Functional titration of each batch of lentivirus was performed in Jurkat, K-562, and THP-1 cells by transducing 4×10^4^ cells via 90 minute spinfection at 1000g and 32°C in 96 well plates with 8µg/mL polybrene (Millipore TR-1003-G). At least five serial dilutions of lentivirus were used, and transductions were performed in quadruplicate. After overnight incubation, media containing lentivirus was removed and replaced with fresh media with and without 2µg/mL puromycin (Gibco A1113803). After five days of selection, cell survival in each well was quantified on a Tecan Spark plate reader using CellTiter-Glo 2.0 Cell Viability Assay (Promega G9242), and percent survival was calculated as the ratio of luminescence in the presence versus absence of puromycin for each lentiviral dilution. Finally, functional lentiviral titer was calculated for all dilutions with 5-30% survival and averaged for each cell line.

### A.4. High-throughput measurements of promoter activity

8×10^7^ Jurkat, K-562, or THP-1 cells were transduced in duplicate via 90 minute spinfection at 1000g and 32°C in 50mL conical tubes with 8×10^6^ infection units (IUs) of virus and 8µg/mL polybrene (Millipore TR-1003-G) for a multiplicity of infection (MOI) of 0.1 and a library coverage of 400X. After transfer to T225 flasks and overnight incubation, media containing lentivirus was removed and replaced with fresh media containing 2µg/mL puromycin (Gibco A1113803). After five days of selection, cells were expanded a further 2-10 days in the absence of puromycin to dilute dead cells and attain at least 4×10^7^ cells (2000X coverage) for sorting. Selected cells were sorted into four 25% bins of EGFP fluorescence using a BD FACSAria Fusion Special Order Research Product. At least 2×10^7^ total cells were sorted for a library coverage of 1000X. Cells from each bin were pelleted, and the supernatant was removed for short-term storage at -20°C.

### A.5. Library preparation and sequencing

Genomic DNA was extracted from sorted cell pellets using Quick-DNA Midiprep Plus Kit (Zymo D4075) using the manufacterer’s instructions. Next generation sequencing (NGS) libraries were prepared using two consecutive PCR steps. In PCR1, the promoters contained in each sorted bin were amplified from the total amount of corresponding genomic DNA using 4µL of each 100µM primer and 400µL NEBNext Ultra II Q5 Master Mix (NEB M0544X). Each 800µL reaction was divided into 8×100µL reactions in a 96 well PCR plate before thermocycling at 98°C for 30s, followed by 20 cycles of 98°C for 10s, 63°C for 30s and 65°C for 45s, with a final extension at 65°C for 5 minutes. All eight completed reactions for each bin were combined into a single tube and vortexed thoroughly before 50µL were purified using a 0.7X AMPure XP bead cleanup (Beckman Coulter A63881). Sequencing adapters and barcodes were then added to the promoter amplicons in PCR2 by combining 2µL of purified PCR1, 2µL of index primers at 10µM each and 25µL NEBNext Ultra II Q5 Master Mix (NEB M0544X). The 50µL reaction was thermocycled at 98°C for 30s, followed by 7 cycles of 98°C for 10s and 65°C for 75s, with a final extension at 65°C for 5 minutes. The PCR2 products were run on a 2% agarose gel, and each produced a single 428bp band, which was extracted using Monarch DNA Gel Extraction Kit (NEB T1020L). Gel-extracted PCR2 products from each bin were then quantified by Qubit 1X dsDNA HS Assay (Thermo Fisher Q33231) and pooled at equimolar ratios before requantification with Qubit and fragment analysis with Agilent 2100 Bioanalyzer using the High Sensitivity DNA Kit (Agilent Technologies 50674626). Prepared libraries were loaded onto the Illumina NextSeq 2000 at 750-850pM and sequenced using 300 cycle v3 kits with P1 or P2 flow cells (Illumina 20050264 and 20046813) to attain at least 1000X sequencing coverage for each replicate.

### A.6. Sequencing analysis

Raw BCL files were converted to fastq files and demultiplexed with bcl-convert v4.0.3 (Illumina). Paired-end reads were trimmed, merged and filtered using fastp (Chen et al., 2018) followed by dereplication and counting with seqfu (Telatin et al., 2021). Only reads with zero mismatches to a promoter in our library were counted, and only promoters with at least five reads in each replicate across all cell lines were considered in downstream analyses.

### A.7. Quantifying expression strength of promoters

The expression strength of each promoter was calculated as the log (base 2) ratio of reads in the highest quartile EGFP bin to the lowest quartile EGFP bin after adding one read to each bin, and the average expression strength (across the two replicates) was calculated for each promoter in each cell line.

## B. Generation of promoter sequences for the experimental dataset

We constructed a promoter library for the experiments described above, which was then used to train and fine-tune our models, containing the following types of sequences.

### B.1. Class I (9991 promoters)

These promoters were extracted from the promoters of endogenous differentially expressed (DE) genes. Gene expression data from LL-100 (Quentmeier et al., 2019) and CCLE (Barretina et al., 2012) were used to identify DE genes. Although we measure expression in Jurkat, K-562 and THP-1 cells, the cell types used for this DE analysis were Jurkat, THP-1 and *NK-92*. We later switched from NK-92s to K-562s due to experimental difficulties. DE genes were identified by DESeq2 (Love et al., 2014). Briefly, for each of the three cell lines, we identified a set of “globally” up/down-regulated genes that were up/down-regulated in that cell line and related cell lines (other immune cells of the same type), when compared to all other cell lines. For each of the three cell lines, we also identified a set of “locally” up/down-regulated genes that were up/down-regulated in that cell line and related cell lines when compared to the other two chosen cell lines and cell lines related to them. For each cell line, we took the intersection of its globally and locally up/down-regulated genes and considered the 1111 top DE genes per cell line (711 up-regulated and 400 down-regulated). Following the rationale from Appendix D.1, we extracted three 250bp promoter sequences for every gene – [TSS - 300bp, TSS - 50bp), [TSS - 100bp, TSS + 150bp), and [TSS + 100bp, TSS + 350bp) – to get a total of 3333 promoters per cell line and 9991 promoters overall (after removing duplicates) to test in our experiments.

### B.2. Class II (7998 promoters)

Promoters in this class were constructed using HOMER (Heinz et al., 2010), a motif detection tool. We supplied the DE genes identified above for the Class I promoters to HOMER, analyzing the [TSS - 300bp, TSS + 50bp] regions of these genes to identify enriched motifs. HOMER identifies two types of enriched motifs, known motifs (which we obtained from Vierstra et al. (2020)) and de-novo motifs. We identified known motifs that were enriched with q-values less than 0.05 and de-novo motifs that were enriched with p-values less than 1e-10. For each cell type, we then generated 2666 promoters, 1500 using motifs enriched in that cell type’s upregulated genes and 1166 using a mix of motifs enriched in that cell type’s upregulated genes and motifs enriched in the other two cell types’ downregulated genes. To generate the promoters, we inserted up to 18 randomly sampled motifs from the above set into an endogenous promoter segment, ([TSS - 100bp, TSS + 150bp)) from an upregulated gene in NK-92s. The exact inserted sequence for each motif was obtained by sampling from its PWM. This process resulted in inserting more than 100bp of motifs into the original 250bp endogenous promoter segment for ∼ 77% of the Class II promoters.

### B.3. Class III (2011 promoters)

Finally, we extracted sequences from the promoters of endogenous highly expressed genes, which were chosen as follows:

1. 1004 genes with the lowest coefficient of variation in their TPM values across all cell lines in the CCLE dataset (restricted to those with a TPM of at least 1).
2. 1007 genes that were up-regulated in all three of the selected cell lines (and related cell lines) vs. all other cell lines in the CCLE dataset, identified using DESeq.

We used the [TSS - 100bp, TSS + 150bp) regions of these genes as 250bp promoter sequences to test in our experiments.

## C. ENCODE accession numbers for the large MPRA dataset we use for evaluation

PE measurements are extracted from element quantification files that derive from count files bearing the following ENCODE accession numbers: ENCFF996ECA, ENCFF018AMJ, ENCFF345ASG, ENCFF970OLE, ENCFF318XMJ, ENCFF821XQZ, ENCFF358MBK, ENCFF379XWL, ENCFF774CHX, ENCFF138DJM, ENCFF277DDE, ENCFF334EKU, ENCFF857FQR, ENCFF259NMG, ENCFF477LDL, ENCFF484JFE, ENCFF227KRF, ENCFF102ZVT, ENCFF418GRL, ENCFF333BAD, ENCFF307HBZ, ENCFF771HPB, ENCFF359KJL, ENCFF035HKU, ENCFF759PPO, ENCFF705AES, ENCFF256WKS, ENCFF352JAC, ENCFF147SMK, ENCFF311DJW, ENCFF350IJA, ENCFF815ORW, ENCFF402GOL, ENCFF865LNO, ENCFF755GRH, ENCFF440YVF, ENCFF703OIL, ENCFF927USI, ENCFF476FXK, ENCFF742ENC, ENCFF112HAT, ENCFF792IHA, ENCFF267VJ. These files were chosen because Gosai et al. (2023)’s lab collected this data and use a subset of these for training PE predictors.

## D. Additional details about pretraining or joint training tasks

### D.1. RNA-seq data

Here, we describe extraction of expression values, and the promoter regions used as inputs to the models during pretraining. TPM values for CCLE and RPKM values from Roadmap are obtained from their respective websites. TPM values for LL-100 are obtained by processing the published raw reads using a standard pipeline (Patel et al., 2022). We filter out any genes that have mean TPM or RPKM values less than 1. Then, we extract three 250bp regions of the promoter for every gene: [TSS - 300bp, TSS - 50bp), [TSS - 100bp, TSS + 150bp), and [TSS + 100bp, TSS + 350bp), which are used to predict expression in every cell line. These regions are chosen by fitting an Xpresso model (Agarwal and Shendure, 2020) to predict median expression across all Roadmap cell lines from various 250bp windows within the TSS ± 1000bp region. We find that the highest prediction performance is obtained using windows within the TSS ± 300bp region. Thus, we choose three 250bp windows covering this region. During training, each promoter sequence window is treated as a separate example with the same associated target expression values. During testing, the predictions for the three windows are averaged to get the final prediction for the gene. We find that this approach yields better fine-tuning and joint training performance compared to using a single large input region such as TSS ± 1500bp.

### D.2. ENCODE TF-binding ChIP-seq data

The process we use to extract peaks is described in this section. We obtain peak calls (narrow peaks) from 1645 TF-binding ChIP-seq datasets from ENCODE that do not have any major quality issues (list of datasets is available in the code repository). Peaks that have a q-value greater than 0.05 are filtered out, and the 1363 cell types with at least 1000 peaks after q-value-based filtering are retained. Because many peaks are very close to each other, we merge peaks that occur within 100bp of each other and create a new unified peak at the mean of the individual peaks’ positions. This unified peak is annotated as being a peak in all datasets from which the individual peaks originated.

### D.3. Summary of datasets

**Table S.1:** Summary of datasets from Section 6.

| Dataset | Assay | Sequence used to train models | Size | Promoter-driven Expression |
| --- | --- | --- | --- | --- |
| LL-100 | RNA-seq of 100 blood cancer cell lines | [TSS - 300 bp, TSS - 50 bp), [TSS - 100 bp, TSS + 150 bp), and [TSS + 100 bp, TSS + 350 bp) | 14,969 | ✗ |
| CCLE | RNA-seq of 1408 cancer cell lines | [TSS - 300 bp, TSS - 50 bp), [TSS - 100 bp, TSS + 150 bp), and [TSS + 100 bp, TSS + 350 bp) | 13,831 | ✗ |
| Roadmap | RNA-seq of 56 cell lines | [TSS - 300 bp, TSS - 50 bp), [TSS - 100 bp, TSS + 150 bp), and [TSS + 100 bp, TSS + 350 bp) | 14,209 | ✗ |
| Sharpr MPRA | MPRA in K-562 and HepG2 | 145 bp sequences whose expression was measured | ~950K | ✓ |
| SuRE MPRA | MPRA in K-562 and HepG2 | 100-500 bp sequences whose expression was measured | ~2.5M (subsampled) | ✓ |
| ENCODE TF-binding ChIP-seq data | 1363 ChIP-seq datasets from diverse cells | +ve set: 600 bp sequence centered at avg position of nearby peaks<br>-ve set: dinucleotide shuffled +ve sequences | ~6M | N/A |

## E. Modelling details

### E.1. Details of the architectures of various models

#### E.1.1. MTLucifer

The MTLucifer model has 3 length-preserving (thus, stride is 1) 1D-convolutional layers followed by 5 transformer layers. The convolutional layers have 256, 512, and 1024 filters of size 5 and use GELU activation (Hendrycks and Gimpel, 2016). After each convolutional layer, we apply a group normalization layer (Wu and He, 2018) with each group comprising 16 channels. The transformer layers have 1024 embedding dimensions, 8 attention heads, and their multi-layer perceptrons have 4096 hidden units. We also use rotary embeddings (Su et al., 2024) with 512 dimensions in each transformer. We apply dropout (Srivastava et al., 2014) to all layers with the dropout probability set to 0.1.

#### E.1.2. Motif occurrences-based FCNs

The smaller motif occurrences-based FCN has 4 layers. We use FIMO (Grant et al., 2011) to extract the number of occurrences of 693 clustered TF-binding motifs (Vierstra et al., 2020) in the sequences (FIMO is run with default arguments and we retain detected motif occurrences with q-value < 0.1). Vectors containing these occurrence counts for all motifs are input to the FCN. Then, 4 fully connected layers with 2048, 1024, 1024 and 512 neurons are applied to get embeddings for each input (each layer except the last uses ReLU activation). These embeddings are then used by a linear output layer to make the PE predictions.

The larger motif occurrences-based FCN is very similar to the smaller one but has 6 layers with 2048, 4096, 4096, 2048, 1024, and 1024 neurons in the 6 layers.

#### E.1.3. CNNs

The smaller CNN has 4 convolutional layers followed by 2 fully connected layers. One-hot encoded sequences are fed as inputs to the network. Then 4 1D length preserving convolutional layers with 512, 768, 768 and 1024 filters of size 5 are applied. Two 1D max pooling layers of size 5 are applied between the second and third layer, and after the last layer. The outputs of the CNN are flattened and passed through 2 fully connected layers with 2048 neurons (and with ReLU activation) and 1024 neurons. The final outputs of this network are then used by a linear output layer to make the PE predictions.

The larger CNN has 6 convolutional layers followed by 2 fully connected layers. The 6 1D length preserving convolutional layers have 512, 768, 768, 1024, 1024, and 1024 filters of size 5. Three 1D max pooling layers of size 5 are applied between the second and third layer, between the fourth and fifth layer, and after the last layer. Again, the outputs of the CNN are flattened and supplied to 2 fully connected layers with 2048 neurons (with ReLU activation) and 1024 neurons. The final outputs are again used by a linear output layer to predict PE.

Note that all convolutional layers in the above CNNs use GELU activation and are followed by group norm (each group has 16 channels) and dropout (0.1 dropout probability) layers.

The ResNet we benchmark has 8 residual blocks. One-hot encoded sequences are first input to a 1D convolutional layer with 512 filters of size 5. Then, 8 standard residual blocks with 512, 512, 768, 768, 1024, 1024, 2048, and 2048 1D filters of size 5 are applied. Adaptive average pooling is performed over the length dimension of the final outputs of the residual blocks and supplied to a fully connected layer with 1024 neurons. Finally, the outputs of this layer are used by a linear output layer to predict PE.

#### E.1.4. LegNets

##### LegNet implementation with the same structure as the model that won the DREAM challenge

The model was constructed using the code provided by Penzar et al. (2022). One-hot encoded sequences are input to the model and the outputs of the final convolutional block are extracted. Adaptive average pooling is then performed over the length dimension of the outputs. Finally, the outputs after pooling are used by a linear output layer to predict PE.

##### Large LegNet with additional filters

We modified the previous LegNet model by increasing the number of filters in the convolutional blocks to 1024, 512, 512, 256, 256, 256, and 256 (from 256, 128, 128, 64, 64, 64, and 64).

#### E.1.5. DNABERT

For fine-tuning DNABERT, we obtain the pretrained model provided by Ji et al. (2021). During fine-tuning, we append a [CLS] token to the tokenized sequences input to the model and use its final embedding to predict PE using a linear output layer.

#### E.1.6. Enformer

We use a Pytorch port^2^ of the original Enformer model published by Avsec et al. (2021) for all experiments. In general, we supply the input sequence to Enformer and get sequence embeddings from its last transformer layer. To aggregate information over the full sequence length, we perform attention pooling and thus get the final sequence embedding. This embedding is then supplied to linear output layers to predict PE.

To perform linear probing of Enformer outputs using Lasso, we first get Enformer outputs from both the human and mouse heads. Then, we flatten these outputs across the length dimension and fit separate Lasso models for each cell type in the PE dataset being modelled using scikit-learn (Pedregosa et al., 2011). The optimal regularization hyperparameter *α* is chosen from the set {1e-5, 1e-4, 1e-3, 1e-2, 1e-1, 1, 10} based on the loss on the validation set. Other hyperparameters are left at their default values.

### E.2. Hyperparameters

1. All models are implemented using PyTorch.
2. We use a cluster consisting of GPU-enabled nodes to train our models. The nodes either use Nvidia TITAN RTXs, A40s or V100s.
3. Unless otherwise stated, all models are trained using the AdawW optimizer (Loshchilov and Hutter, 2017).
4. For regression-based pretraining tasks from Section 6, we Z-score all target values before fitting models.
5. When training from scratch to model the fluorescence data, we generally use a 1e-5 learning rate, 1e-4 weight decay and 96 batch size. Models are trained for a maximum of 50 epochs but if the average Spearman’s rank correlation coefficient of the validation set does not improve for 5 epochs, we stop training. We change our hyperparameters in the following cases:
  - When training DNABERT from scratch (i.e. random initialization), we use a batch size of 64 due to GPU memory constraints.
  - While training LegNets, we employ the optimizer (AdamW) and scheduler (One Cycle Learning Rate Policy (Smith and Topin, 2019)) used by Penzar et al. (2022) to train LegNet models for the DREAM challenge. For the smaller LegNet, we use 0.005 learning rate, 0.01 weight decay and 1024 batch size. For the larger one, we use 0.005 learning rate, 0.01 weight decay and 192 batch size. The scheduler uses the same hyperparameters as those used by Penzar et al. (2022).
  - When training MPRAnn, we employ the optimizer (Adam) and hyperparameters used by Agarwal et al. (2023) i.e. 0.001 learning rate and batch size of 32.
  - When training Malinois, we employ the optimizer (AdamW) and scheduler (cosine annealing witg warm restarts) used by Gosai et al. (2023). As prescribed by them, we use a 0.0032658700881052086 learning rate and 0.0003438210249762151 weight decay. We use a batch size of 512 and train for a maximum of 200 epochs but if the average Spearman’s rank correlation coefficient of the validation set does not improve for 30 epochs, we stop training.
  - When training Enformer from scratch (i.e. random initialization), we use a 5e-4 learning rate, 5e-4 weight decay and 96 batch size. We note that using a higher learning rate led to training instability.
6. When training from scratch to model the Malinois MPRA data, we generally use a 1e-4 learning rate, 1e-4 weight decay and 96 batch size. Models are trained for a maximum of 50 epochs but if the average Spearman’s rank correlation coefficient of the validation set does not improve for 5 epochs, we stop training. We change our hyperparameters in the following cases:
  - When training DNABERT from scratch (i.e. random initialization), we use a 1e-5 learning rate and a batch size of 64 due to GPU memory constraints.
  - While training LegNets, we employ the optimizer (AdamW) and scheduler (One Cycle Learning Rate Policy (Smith and Topin, 2019)) used by Penzar et al. (2022) to train LegNet models for the DREAM challenge. For the smaller LegNet, we use 0.05 learning rate, 0.01 weight decay and 1024 batch size. For the larger one, we use 0.01 learning rate, 0.01 weight decay and 192 batch size. The scheduler uses the same hyperparameters as those used by Penzar et al. (2022).
  - When training MPRAnn, we employ the optimizer (Adam) and hyperparameters used by Agarwal et al. (2023) i.e. 0.001 learning rate and batch size of 32.
  - When training Malinois, we employ the optimizer (AdamW) and scheduler (cosine annealing witg warm restarts) used by Gosai et al. (2023). As prescribed by them, we use a 0.0032658700881052086 learning rate, 0.0003438210249762151 weight decay and a batch size of 1076. We train for a maximum of 200 epochs but if the average Spearman’s rank correlation coefficient of the validation set does not improve for 30 epochs, we stop training.
7. When pretraining MTLucifer on the tasks from Section 6, we use a 1e-5 learning rate and 1e-4 weight decay. We train for 50 epochs and stop training if the overall validation loss does not improve for 5 epochs. We use the following batch sizes due to GPU memory constraints:
  - RNA-seq: 96
  - ENCODE TF-binding ChIP-seq: 32
  - Sharpr-MPRA: 96
  - SuRE: 24
  - SuRE, Sharpr-MPRA: 24
8. When we fine-tune the pretrained MTLucifer models on the fluorescence data, we use a 1e-5 learning rate, 1e-4 weight decay and a batch size of 96. Models are trained for a maximum of 50 epochs but if the average Spearman’s rank correlation coefficient of the validation set does not improve for 5 epochs, we stop training.
9. When we perform linear probing of the pretrained MTLucifer models on the fluorescence data, we use a 1e-3 learning rate, 1e-4 weight decay and a batch size of 96. Models are trained for a maximum of 50 epochs but if the average Spearman’s rank correlation coefficient of the validation set does not improve for 5 epochs, we stop training.
10. When jointly training MTLucifer on the tasks from Section 6 and the fluorescence dataset, we use a 1e-5 learning rate and 1e-4 weight decay. We train for 50 epochs and stop training if the average Spearman’s rank correlation coefficient of the fluorescence dataset’s validation set does not improve for 5 epochs. We use the following batch sizes due to GPU memory constraints:
  - RNA-seq: 32
  - ENCODE TF-binding ChIP-seq: 32
  - Sharpr-MPRA: 64
  - SuRE: 12
  - SuRE, Sharpr-MPRA: 12
11. When we fine-tune pretrained DNABERT on the fluorescence data, we use a 1e-5 learning rate, 1e-4 weight decay and a batch size of 64. Models are trained for a maximum of 50 epochs but if the average Spearman’s rank correlation coefficient of the validation set does not improve for 5 epochs, we stop training.
12. When we fine-tune pretrained Enformer on the fluorescence data, we use a 1e-3 learning rate, 5e-4 weight decay and a batch size of 96. Models are trained for a maximum of 50 epochs but if the average Spearman’s rank correlation coefficient of the validation set does not improve for 5 epochs, we stop training.
13. When we fine-tune the pretrained MTLucifer models on the Malinois MPRA data, we use a 1e-4 learning rate, 1e-4 weight decay and a batch size of 96. Models are trained for a maximum of 50 epochs but if the average Spearman’s rank correlation coefficient of the validation set does not improve for 5 epochs, we stop training.
14. When we perform linear probing of the pretrained MTLucifer models on the fluorescence data, we use a 1e-3 learning rate, 1e-4 weight decay and a batch size of 96. Models are trained for a maximum of 50 epochs but if the average Spearman’s rank correlation coefficient of the validation set does not improve for 5 epochs, we stop training.
15. When jointly training MTLucifer on the tasks from Section 6 and the Malinois MPRA dataset, we use a 1e-4 learning rate and 1e-4 weight decay. We train for 50 epochs and stop training if the average Spearman’s rank correlation coefficient of the fluorescence dataset’s validation set does not improve for 5 epochs. We use the following batch sizes due to GPU memory constraints:
  - RNA-seq: 32
  - ENCODE TF-binding ChIP-seq: 32
  - Sharpr-MPRA: 64
  - SuRE: 8
  - SuRE, Sharpr-MPRA: 8
16. When we fine-tune pretrained DNABERT on the Malinois MPRA data, we use a 1e-5 learning rate, 1e-4 weight decay and a batch size of 64. Models are trained for a maximum of 50 epochs but if the average Spearman’s rank correlation coefficient of the validation set does not improve for 5 epochs, we stop training.
17. When we fine-tune pretrained Enformer on the Malinois MPRA data, we use a 1e-4 learning rate, 1e-4 weight decay and a batch size of 96. Models are trained for a maximum of 50 epochs but if the average Spearman’s rank correlation coefficient of the validation set does not improve for 5 epochs, we stop training.

## F. Benchmarking using subsampled Malinois MPRA data

We create a smaller subsampled Malinois MPRA dataset by subsampling the training set to 15,000 examples (with expression measurements in all 5 cell types) to roughly match the size of the fluorescence dataset while keeping the full validation and test sets. The subsampling is performed while keeping the proportion of sequences from each chromosome in the train set constant. The performances of various model architectures when trained from scratch are presented in Table S.2 and the performances of various transfer learning approaches are presented in Table S.3. We see that MTLucifer is still the best architecture when trained from scratch. When we use transfer learning, we see a substantial improvement in performance - fine-tuning Enformer improves performance by 26-51% when compared to the best model that was trained from scratch. When using the fluorescence dataset, this approach was only slightly worse than performing linear probing on Enformer predictions using a Lasso regression model. We also observe that pretraining MTLucifer on the MPRA data before fine-tuning it is the second best approach. In conclusion, these trends are very similar to those we observe when using the small fluorescence dataset, showing that the size of the training dataset is the main determinant of relative performances.

**Table S.2:** Prediction performance obtained using various model architectures when trained from scratch on the subsampled Malinois MPRA dataset.

| Model Class | Number of parameters | HepG2 |  | K-562 |  | SK-N-SH |  | A549 |  | GM12878 |  |
| --- | --- | --- | --- | --- | --- | --- | --- | --- | --- | --- | --- |
| | | $r$ | $\rho$ | $r$ | $\rho$ | $r$ | $\rho$ | $r$ | $\rho$ | $r$ | $\rho$ |
| MTLucifer | 66.3M | **0.4876** | **0.4948** | 0.4715 | **0.4613** | **0.5044** | **0.5644** | 0.4308 | 0.4089 | **0.2883** | 0.2883 |
| Motif-based FCN | 5.1M | 0.2930 | 0.2454 | 0.2834 | 0.2046 | 0.2826 | 0.2611 | 0.2650 | 0.2299 | 0.1675 | 0.1724 |
| Large motif-based FCN | 38.1M | 0.2984 | 0.2433 | 0.2925 | 0.2066 | 0.2862 | 0.2601 | 0.2703 | 0.2265 | 0.1726 | 0.1721 |
| CNN | 11.0M | 0.4016 | 0.3998 | 0.3769 | 0.3577 | 0.4236 | 0.4659 | 0.3419 | 0.3253 | 0.2212 | 0.2308 |
| Large CNN | 21.5M | 0.3897 | 0.3797 | 0.3592 | 0.3107 | 0.4020 | 0.4241 | 0.3302 | 0.2830 | 0.2171 | 0.2204 |
| ResNet | 114M | 0.3719 | 0.3577 | 0.3415 | 0.3052 | 0.3693 | 0.3978 | 0.2993 | 0.2769 | 0.1870 | 0.2187 |
| LegNet | 1.8M | 0.4729 | 0.4426 | **0.4766** | 0.4219 | 0.4762 | 0.5025 | 0.4205 | 0.3912 | 0.2064 | 0.2121 |
| Large LegNet | 33.1M | 0.4817 | 0.4456 | 0.5122 | 0.4538 | 0.4871 | 0.5184 | **0.4868** | **0.4382** | 0.2716 | **0.3012** |
| MPRAnn | 808K | 0.3526 | 0.3624 | 0.3316 | 0.3175 | 0.3495 | 0.4039 | 0.2939 | 0.3105 | 0.1493 | 0.1716 |
| Malinois | 4.5M | 0.3337 | 0.3471 | 0.3227 | 0.3204 | 0.3650 | 0.3943 | 0.3116 | 0.2809 | 0.1921 | 0.1836 |
| DNABERT (random initialization) | 89.2M | 0.4458 | 0.4272 | 0.4323 | 0.4007 | 0.4556 | 0.4675 | 0.3988 | 0.3660 | 0.2268 | 0.2474 |
| Enformer (random initialization) | 229M | 0.3709 | 0.2616 | 0.3846 | 0.2877 | 0.3437 | 0.2141 | 0.3710 | 0.3558 | 0.2024 | 0.2315 |

**Table S.3:** Prediction performances obtained using various training strategies when used to model the subsampled Malinois MPRA dataset.

| Model Class | Pretraining or Joint Training Tasks | Transfer Method | HepG2 |  | K-562 |  | SK-N-SH |  | A549 |  | GM12878 |  |
| --- | --- | --- | --- | --- | --- | --- | --- | --- | --- | --- | --- | --- |
| | | | $r$ | $\rho$ | $r$ | $\rho$ | $r$ | $\rho$ | $r$ | $\rho$ | $r$ | $\rho$ |
| MTLucifer | - | Training from scratch | 0.4876 | 0.4948 | 0.4715 | 0.4613 | 0.5044 | 0.5644 | 0.4308 | 0.4089 | 0.2883 | 0.2883 |
| MTLucifer | All RNA-Seq | Joint training | 0.4802 | 0.4790 | 0.4577 | 0.4346 | 0.5026 | 0.5584 | 0.4355 | 0.4167 | 0.2857 | 0.2732 |
| MTLucifer | ENCODE TF-binding ChIP-seq | Joint training | 0.4982 | 0.4961 | 0.4949 | 0.4561 | 0.5037 | 0.5500 | 0.4509 | 0.4377 | 0.2857 | 0.2955 |
| MTLucifer | Sharpr-MPRA | Joint training | 0.4896 | 0.4912 | 0.4683 | 0.4455 | 0.5050 | 0.5569 | 0.4374 | 0.4141 | 0.2724 | 0.2709 |
| MTLucifer | SuRE MPRA | Joint training | 0.5756 | 0.5435 | 0.5993 | 0.5250 | 0.5870 | 0.5996 | 0.5618 | 0.4933 | 0.3499 | 0.3403 |
| MTLucifer | Sharpr, SuRE MPRA | Joint training | 0.5581 | 0.5334 | 0.5644 | 0.4998 | 0.5683 | 0.5891 | 0.5290 | 0.4909 | 0.3141 | 0.3169 |
| MTLucifer | All RNA-Seq | Fine-tuning | 0.4854 | 0.4832 | 0.4809 | 0.4555 | 0.5083 | 0.5562 | 0.4429 | 0.4167 | 0.2936 | 0.2860 |
| MTLucifer | ENCODE TF-binding ChIP-seq | Fine-tuning | 0.5350 | 0.5255 | 0.5451 | 0.4943 | 0.5488 | 0.5833 | 0.5078 | 0.4783 | 0.3073 | 0.3055 |
| MTLucifer | Sharpr-MPRA | Fine-tuning | 0.5398 | 0.4984 | 0.5344 | 0.4541 | 0.5410 | 0.5520 | 0.5093 | 0.4542 | 0.2975 | 0.2927 |
| MTLucifer | SuRE MPRA | Fine-tuning | 0.6302 | 0.6005 | 0.6460 | 0.5867 | 0.6203 | 0.6307 | 0.5938 | 0.5352 | 0.3698 | 0.3569 |
| MTLucifer | Sharpr, SuRE MPRA | Fine-tuning | 0.6450 | 0.6098 | 0.6652 | 0.5973 | 0.6419 | 0.6464 | 0.6123 | 0.5567 | 0.3781 | 0.3680 |
| MTLucifer | All RNA-Seq | Linear probing | 0.3919 | 0.4001 | 0.3652 | 0.3491 | 0.4199 | 0.4782 | 0.3359 | 0.3460 | 0.2252 | 0.2467 |
| MTLucifer | ENCODE TF-binding ChIP-seq | Linear probing | 0.4217 | 0.4071 | 0.4183 | 0.3726 | 0.4328 | 0.4454 | 0.3878 | 0.3737 | 0.2476 | 0.2726 |
| MTLucifer | Sharpr-MPRA | Linear probing | 0.5111 | 0.4658 | 0.4940 | 0.4106 | 0.5057 | 0.5064 | 0.4546 | 0.4045 | 0.2616 | 0.2583 |
| MTLucifer | SuRE MPRA | Linear probing | 0.6179 | 0.5657 | 0.6480 | 0.5579 | 0.6102 | 0.5899 | 0.5907 | 0.4919 | 0.3549 | 0.3167 |
| MTLucifer | Sharpr, SuRE MPRA | Linear probing | 0.6278 | 0.5879 | 0.6491 | 0.5756 | 0.6181 | 0.6144 | 0.5897 | 0.5220 | 0.3538 | 0.3408 |
| DNABERT | Human Genome MLM | Fine-tuning | 0.3625 | 0.3421 | 0.3309 | 0.2814 | 0.3783 | 0.3843 | 0.3158 | 0.2791 | 0.1962 | 0.1899 |
| Enformer | Variety of genomic and epigenomic data | Fine-tuning | **0.7274** | **0.6888** | **0.7622** | **0.6871** | **0.7095** | **0.7103** | **0.6714** | **0.6192** | **0.4094** | **0.4347** |
| Enformer | Variety of genomic and epigenomic data | Linear probing with Lasso | 0.5973 | 0.5369 | 0.6261 | 0.4960 | 0.6065 | 0.5803 | 0.5771 | 0.4778 | 0.3411 | 0.3226 |
| Mean increase in performance of best-performing method vs. training from scratch |  |  | 49.18% | 39.21% | 61.65% | 48.95% | 40.66% | 25.85% | 55.85% | 51.43% | 42.00% | 50.78% |

## Footnotes

1 https://dreamchallenges.org/predicting-gene-expression/

2 https://github.com/lucidrains/enformer-pytorch

## Notes

https://github.com/anikethjr/promoter_models

### Competing Interest Statement

The authors have declared no competing interest.

### Summary of Updates

More comprehensive benchmarking is performed in this version of the manuscript. We benchmark more architectures and transfer learning methods, and the benchmarking is now performed in two settings - a data-constrained setting and a large dataset setting.

